# Machine Learning Prediction of Antimicrobial Response in *Pleurotus ostreatus* Extracts Cultivated on Cassava Peel: A Proof-of-Concept Study

**DOI:** 10.64898/2026.08.10.743970

**Authors:** Olagunju Johnson Adetuwo

## Abstract

Antimicrobial resistance has intensified the search for sustainable natural products with antimicrobial properties. *Pleurotus ostreatus* cultivated on lignocellulosic agro-wastes, including cassava peel, offers potential for bioactive-compound production and agricultural waste valorization. Conventional antimicrobial screening, however, can be labour-intensive when multiple extracts and pathogens are evaluated. This study evaluated whether extraction solvent, broad pathogen taxonomic category, and batch-level mycochemical composition could predict the antimicrobial response of *P. ostreatus* extracts cultivated on cassava peel and identified the variables contributing most strongly to prediction. Ethanolic and aqueous mushroom extracts were evaluated against seven microbial pathogens using agar well diffusion and broth microdilution assays. The dataset comprised 42 observations. A Random Forest model with leave-one-out cross-validation (LOOCV) was used to model zone of inhibition as a regression task and minimum inhibitory concentration (MIC) as a binary classification task. The Random Forest regression model showed moderate internal predictive performance for zone of inhibition (R^2^ = 0.68, MAE = 0.62 mm, RMSE = 0.75 mm). Extraction solvent was the strongest predictor, whereas batch-level mycochemical variables contributed minimally. In contrast, MIC classification performed poorly (accuracy = 0.43; F1-score = 0.33), indicating that the available predictors were insufficient to discriminate the two observed MIC groups. The findings support machine learning as an exploratory complement to antimicrobial screening of mushroom-derived natural products. Given the limited dataset and three cultivation batches, the results are preliminary. Larger, multi-substrate and multi-species datasets with replicate-resolved biochemical measurements will be required to develop robust predictive models.

## 1. Introduction

Antimicrobial resistance (AMR) represents one of the most significant global public health challenges of the twenty-first century. The rapid emergence of multidrug-resistant bacterial and fungal pathogens has reduced the effectiveness of many conventional antimicrobial agents, creating an urgent need for alternative therapeutic compounds from natural sources. Edible mushrooms have attracted increasing scientific attention because they synthesize diverse secondary metabolites with antimicrobial, antioxidant, immunomodulatory, and anti-inflammatory properties (Ferreira *et al*., 2009; Fakoya *et al*., 2020; Elhusseiny *et al*., 2021; Albertó, 2021; Adedokun *et al*., 2022)

Among cultivated edible mushrooms, *Pleurotus ostreatus* (oyster mushroom) is particularly valued because of its nutritional quality, ease of cultivation, environmental adaptability, and ability to grow efficiently on inexpensive lignocellulosic agricultural residues. These characteristics make *P. ostreatus* an attractive candidate for sustainable biotechnology, simultaneously supporting food production and agricultural waste management (Carrasco-González *et al*., 2017; Gashaw, 2020; Adetuwo et al., 2026).

Cassava peel constitutes one of the most abundant agricultural residues generated in tropical countries. Although frequently discarded as waste, cassava peel contains cellulose, hemicellulose, and lignin that provide suitable substrates for mushroom cultivation. Its utilization therefore contributes to circular bioeconomy principles by converting low-value agricultural waste into value-added food products and biologically active compounds.

Previous investigations have demonstrated that *P. ostreatus* extracts cultivated on diverse agricultural wastes possess antimicrobial activity against a broad spectrum of Gram-positive bacteria, Gram-negative bacteria, and opportunistic fungi (Gashaw *et al*., 2020). These biological activities have been attributed to phenolic compounds, flavonoids, terpenoids, alkaloids, steroids, tannins, and other secondary metabolites extracted from mushroom fruiting bodies. However, evaluating antimicrobial activity experimentally requires extensive laboratory testing involving multiple extraction methods, microbial species, biological replicates, and susceptibility assays, making large-scale screening expensive and time-consuming.

Machine learning (ML) has emerged as a valuable computational approach for analysing complex biological datasets and identifying predictive relationships among experimental variables. In biomedical and pharmaceutical research, ML algorithms have been applied to drug discovery, antimicrobial peptide identification and design, metabolomics, microbial genomics, and natural product screening (Yuan et al., 2023; Arnold et al., 2023; Prihoda et al., 2021; Melo et al., 2021; Wan et al., 2024). Beyond natural product discovery, ML has also been applied directly to antimicrobial resistance prediction, with systematic reviews describing the use of models including Random Forest for antimicrobial susceptibility classification (Tang et al., 2022; Sakagianni et al., 2023; Lv & Wang, 2024; Li et al., 2024). Nevertheless, the application of ML to antimicrobial datasets generated from edible mushrooms cultivated on agricultural wastes remains limited.

Random Forest algorithms can model nonlinear relationships, accommodate mixed categorical and continuous variables, and provide estimates of predictor importance, making them useful for exploratory biological datasets (Ghosh & Cabrera, 2022). When combined with leave-one-out cross-validation (LOOCV), Random Forest provides a practical framework for estimating internal predictive performance when conventional train-test partitioning is impractical because of limited sample size.

The present study therefore investigated whether experimental characteristics and broad pathogen categories rather than pathogen identity itself could predict antimicrobial response in *P. ostreatus* extracts cultivated on cassava peel, using experimental antimicrobial assay data generated from aqueous and ethanolic extracts. This framing was deliberately chosen over the broader claim that the model can predict antimicrobial potential in general, since pathogen identity was intentionally excluded as a predictor (Section 2.3). Specifically, the study aimed to (i) predict zone of inhibition using regression analysis, (ii) classify minimum inhibitory concentration (MIC) outcomes, and (iii) identify the experimental variables that contribute most strongly to antimicrobial performance. Given the relatively small dataset, the analysis is presented as a proof-of-concept methodological demonstration rather than a fully validated predictive model, providing a foundation for future studies involving larger multi-substrate and multi-species datasets.

## 2. Materials and Methods

### 2.1 Study Design

This study employed a proof-of-concept machine learning approach to investigate the feasibility of predicting the antimicrobial activity of *Pleurotus ostreatus* cultivated on cassava peel. Rather than replacing conventional microbiological assays, the computational model was designed to complement laboratory screening by identifying the experimental variables that most strongly influenced antimicrobial performance. Owing to the relatively small dataset, the investigation was intended as an exploratory demonstration of machine learning applied to mushroom-derived antimicrobial data.

### 2.2 Experimental Dataset

The dataset used for machine learning analysis was generated from antimicrobial susceptibility experiments involving *P. ostreatus* cultivated exclusively on cassava peel.

Ethanolic and aqueous extracts of the mushroom fruiting bodies were evaluated against seven clinically important microbial pathogens:

- *Escherichia coli*
- *Staphylococcus aureus*
- *Klebsiella pneumoniae*
- *Pseudomonas aeruginosa*
- *Salmonella typhi*
- *Candida albicans*
- *Aspergillus fumigatus*

Antimicrobial activity was determined using two complementary laboratory techniques:

- Agar well diffusion assay to measure zone of inhibition (ZOI, mm).
- Broth microdilution assay to determine minimum inhibitory concentration (MIC, µg/mL).

Each antimicrobial assay was performed in triplicate for both extraction solvents, producing a total of 42 observations (7 pathogens × 2 extract types × 3 biological replicates). Each observation was linked to the corresponding batch-level mycochemical profile, including phenolics, flavonoids, tannins, terpenoids, alkaloids, steroids, and saponins, which served as continuous predictor variables.

### 2.3 Predictor Variables

The predictor variables incorporated into the machine learning models comprised both categorical and continuous variables. Categorical variables included extraction solvent (ethanolic or aqueous) and pathogen taxonomic class (Gram-positive bacteria, Gram-negative bacteria, or fungi). Continuous variables consisted of the measured concentrations of the seven mycochemical constituents obtained from each cultivation and extraction batch.

To minimize overfitting and prevent species-level memorization, individual pathogen identity was intentionally excluded from model development. Instead, pathogens were grouped according to broader taxonomic classifications, thereby encouraging the model to identify generalizable predictor–response associations rather than memorize organism-specific responses. This choice narrows the scope of the research question: rather than predicting antimicrobial potential for a specific named pathogen, the model addresses whether experimental characteristics and broad pathogen categories can predict antimicrobial response in *P. ostreatus* extracts. Given the study design, inclusion of pathogen identity could also dominate prediction and mask the influence of the experimental variables of primary interest.

### 2.4 Machine Learning Model Development

A Random Forest algorithm (Breiman, 2001) was selected because it can accommodate both categorical and continuous predictors and capture nonlinear relationships while providing estimates of predictor importance. The model was implemented in Python using the scikit-learn library (Pedregosa et al., 2011), with 300 decision trees (estimators) and a maximum tree depth of 4. These parameters were selected to balance predictive performance with model complexity and to limit overfitting within the available dataset.

### 2.5 Cross-Validation Strategy

Given the limited sample size, conventional train-test data partitioning would have substantially reduced the amount of information available for model training. Consequently, leave-one-out cross-validation (LOOCV) was adopted as the primary evaluation strategy. Under LOOCV, one observation was excluded during each iteration for model testing, while the remaining observations were used for training. This process was repeated until every observation had served once as the validation sample, providing an internal estimate of predictive performance appropriate for a small dataset, although not a substitute for independent validation.

Because the batch-level mycochemical predictors (Section 2.2) are shared across the 14 observations drawn from each cultivation batch, standard LOOCV allows observations from the same batch to appear in both the training and validation folds within a given iteration. This could, in principle, allow the model to exploit batch identity rather than generalizable biological relationships. To assess this risk directly, leave-one-batch-out cross-validation was additionally performed, in which all observations from a given cultivation batch were held out together (three folds corresponding to the three cultivation replicates). If batch-level leakage were materially inflating the standard LOOCV estimates, leave-one-batch-out performance would be expected to be substantially lower. This was not observed (Section 3.1), which is consistent with the categorical predictors (extract type and organism class), which were balanced across the three batches, driving model predictions rather than the batch-specific mycochemical values, whose feature importance was minimal.

### 2.6 Prediction Tasks

Two independent prediction tasks were performed. Zone of inhibition (mm) was treated as a continuous response variable and analysed using Random Forest regression, with performance assessed using the coefficient of determination (R^2^), mean absolute error (MAE), and root mean square error (RMSE).

The MIC dataset contained only two observed values (25 and 50 µg/mL); no predefined clinical breakpoint was applied. Consequently, MIC prediction was reformulated as a binary classification problem defined directly by these two observed values: “high sensitivity” (MIC = 25 µg/mL; 24 of 42 observations—E. coli, S. typhi, C. albicans, and A. fumigatus) and “low sensitivity” (MIC = 50 µg/mL; 18 of 42 observations—S. aureus, K. pneumoniae, and P. aeruginosa). Classification was evaluated using LOOCV accuracy, F1-score, and a confusion matrix. This strategy was adopted because binary classification was more appropriate than regression for the two-valued MIC outcomes actually observed; it should be interpreted as a data-driven grouping of observed responses rather than a clinically validated susceptibility threshold.

### 2.7 Feature Importance Analysis

To identify the variables contributing most strongly to antimicrobial prediction, feature importance values generated by the Random Forest algorithm were examined. Because categorical predictors (extract type and pathogen taxonomic class) were one-hot encoded, the Random Forest importance algorithm assigns a separate importance value to each encoded level (e.g., “ethanolic” and “aqueous” as distinct columns). These values were summed within each original variable to report a single combined importance per predictor and avoid the interpretive ambiguity of treating encoding artefacts as independent findings. Per-level importance values are additionally provided in the deposited results file (Adetuwo, 2026) for transparency.

### 2.8 Study Limitations

The machine learning analysis was intentionally presented as an exploratory proof-of-concept because the dataset comprised only 42 observations obtained from three cultivation replicates. Furthermore, biochemical measurements were collected at the batch level rather than for each antimicrobial replicate, thereby limiting the resolution of the predictor variables. Finally, the model was developed exclusively using *P*.*ostreatus* cultivated on cassava peel; therefore, its predictive performance cannot yet be generalized to other mushroom species or cultivation substrates.

## 3. Results

### 3.1 Prediction of Zone of Inhibition

The Random Forest regression model demonstrated moderate internal predictive performance for estimating the zone of inhibition (ZOI) of *Pleurotus ostreatus* extracts. Using LOOCV, the model achieved a coefficient of determination (R^2^) of 0.68, a mean absolute error (MAE) of 0.62 mm, and a root mean square error (RMSE) of 0.75 mm. These results indicate that the selected predictor variables accounted for a substantial proportion of the observed variability in antimicrobial activity despite the limited dataset.

The observed level of predictive performance suggests that meaningful predictor–response associations existed among extraction solvent, pathogen taxonomic class, and antimicrobial response. Nevertheless, because the model was developed using a relatively small number of observations, these performance metrics should be interpreted as preliminary internal predictive associations rather than evidence of a fully validated, generalizable predictive model.

**Figure 1.**
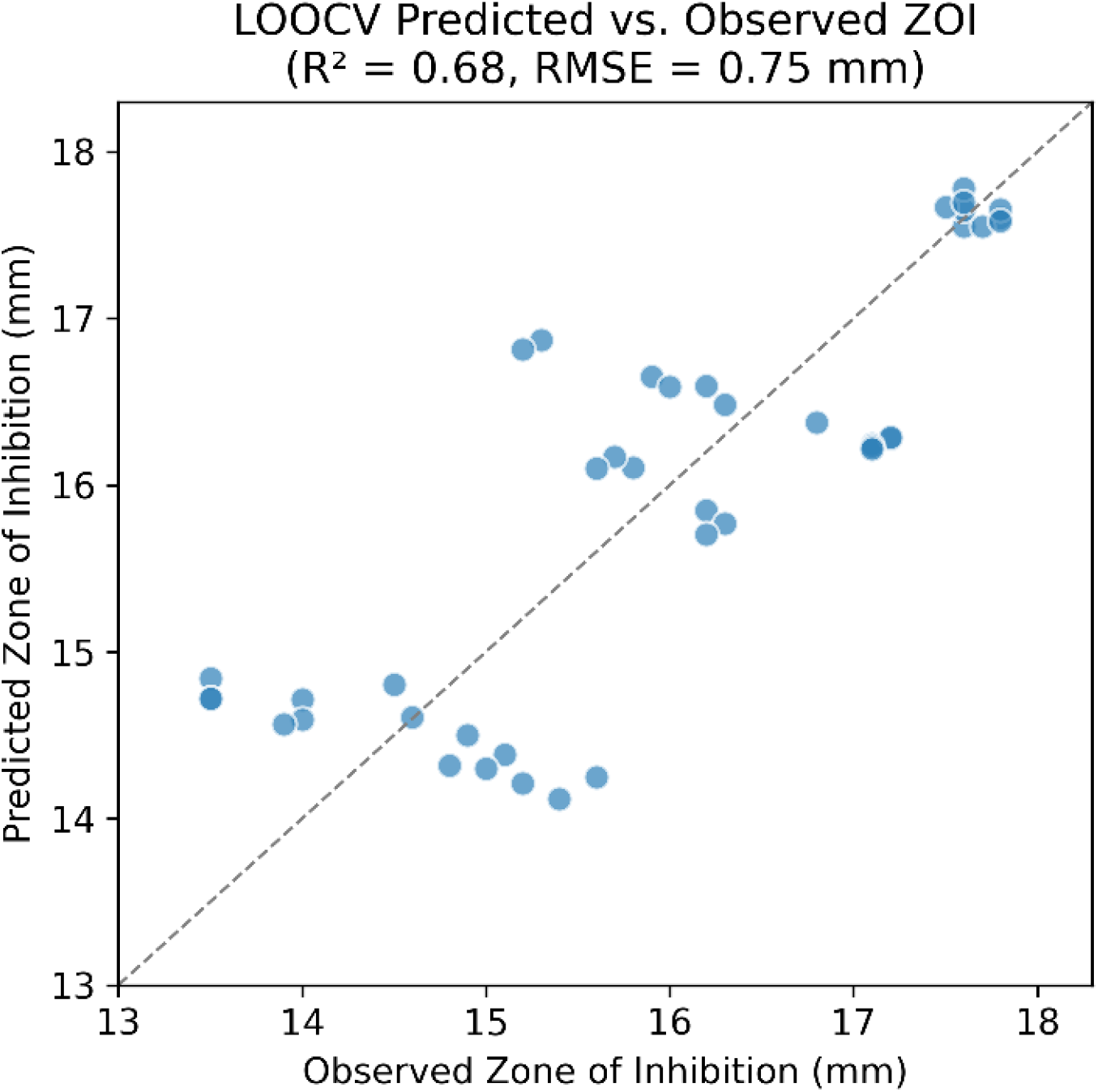
Leave-one-out cross-validated predicted versus observed zone of inhibition (mm).

### 3.2 Feature Importance Analysis

Feature importance analysis identified extract type (ethanolic versus aqueous; combined importance = 69.6%) as the dominant predictor of zone of inhibition. Pathogen taxonomic class contributed a smaller but non-trivial combined importance (27.1%), whereas the seven batch-level mycochemical variables together contributed 3.3% of total model importance.

The minimal combined contribution of the biochemical variables is likely attributable to the limited variation observed among the three cultivation batches rather than the absence of biological relevance. As reported in Section 2.5, leave-one-batch-out cross-validation produced performance at least as strong as standard LOOCV (R^2^ = 0.81 versus 0.68). This result provides no indication that the observed LOOCV performance was materially inflated by batch-level leakage. Future investigations involving a greater number of cultivation batches and more diverse substrate compositions may reveal stronger relationships between mycochemical composition and antimicrobial activity.

**Figure 2.**
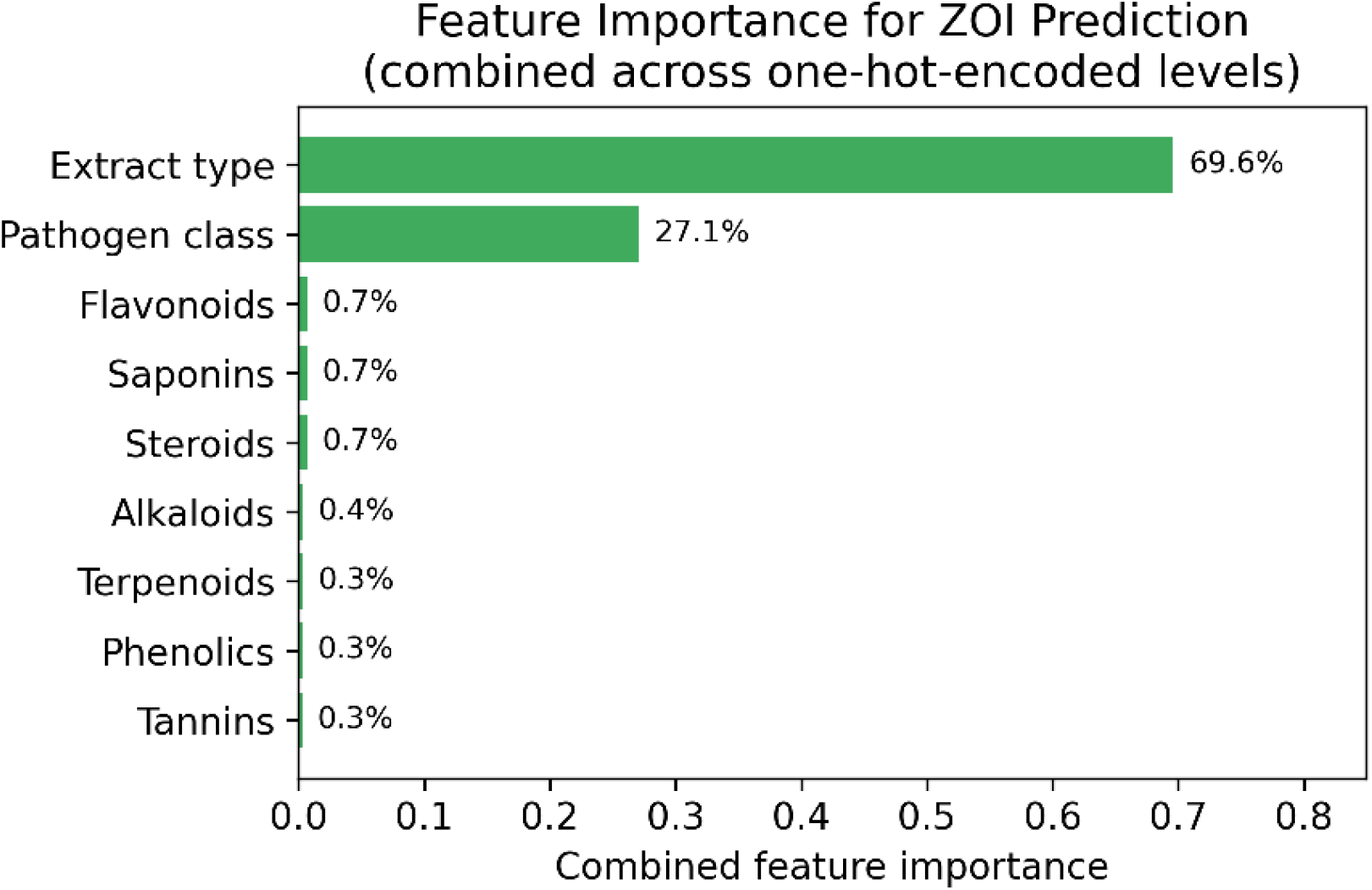
Feature importance for zone-of-inhibition prediction, combined across one-hot-encoded levels within each original categorical variable (extract type and pathogen class).

### 3.3 Prediction of Minimum Inhibitory Concentration

Unlike the regression model for zone of inhibition, the Random Forest classifier demonstrated poor performance when predicting the two MIC sensitivity groups. Under LOOCV, the classifier achieved an accuracy of 0.43 and an F1-score of 0.33, both indicating limited discriminatory performance and an accuracy below the majority-class baseline of 0.57.

The reduced predictive performance indicates that the selected predictor variables were insufficient to discriminate reliably between the two observed MIC groups. In particular, differences among Gram-negative bacterial species resulted in inconsistent susceptibility patterns that could not be adequately captured using pathogen taxonomic class alone, a pattern consistent with the established role of species-specific outer membrane permeability and efflux pump activity in shaping intrinsic resistance among Gram-negative organisms (Nikaido, 1998).

**Figure 3.**
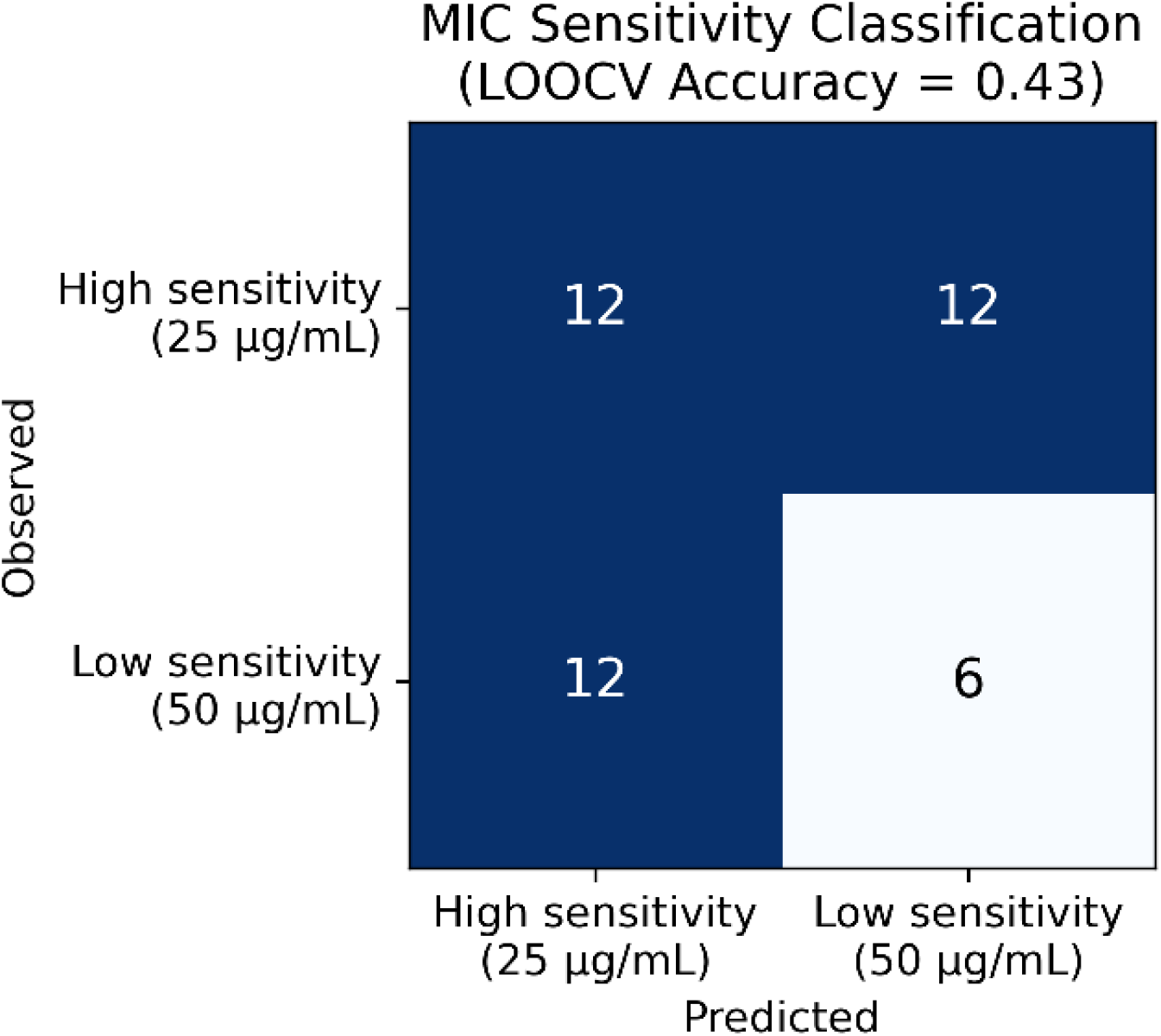
Confusion matrix for prediction of the two data-driven MIC sensitivity groups.

### 3.4 Summary of Model Performance

Overall, the Random Forest regression model demonstrated reasonable internal predictive capability for zone of inhibition, whereas the classification model performed poorly for the two observed MIC groups. These findings indicate that the available predictor set was more informative for modelling the continuous ZOI response than for discriminating the limited binary MIC outcomes in this dataset.

| Target | Task | Key metric(s) | LOOCV performance |
| --- | --- | --- | --- |
| Zone of inhibition (mm) | Regression | R<sup>2</sup> / MAE / RMSE | 0.68 / 0.62 mm / 0.75 mm |
| MIC sensitivity group | Classification | Accuracy / F1 | 0.43 / 0.33 (baseline = 0.57) |

## 4. Discussion

The present study explored the application of a Random Forest machine learning algorithm to predict the antimicrobial activity of *Pleurotus ostreatus* cultivated on cassava peel. Although exploratory in nature, the findings show how computational modelling can complement conventional microbiological screening by identifying experimental variables associated with antimicrobial response. The regression model showed moderate internal predictive performance despite the relatively small dataset.

A key finding was that extraction solvent emerged as the dominant predictor of zone of inhibition. This observation is biologically plausible because ethanol can extract a broader range of moderately polar and non-polar bioactive compounds than water. The stronger antimicrobial response associated with ethanolic extracts is therefore consistent with differences in the solubility of phenolic compounds, flavonoids, terpenoids, and related secondary metabolites in organic solvents, and aligns with the stronger influence of extract polarity reported in the parent substrate-comparison study on which this dataset is based (Adetuwo et al., 2026). Importantly, the model identifies extraction solvent as the strongest predictor; it does not by itself establish a causal effect of solvent polarity.

The relatively small contribution of individual mycochemical variables should not be interpreted as evidence that these compounds lack antimicrobial significance. Rather, the limited variation in biochemical composition across only three cultivation replicates constrained the model’s ability to distinguish their individual contributions (Adetuwo et al., 2026). This limitation highlights the importance of replicate-resolved biochemical measurements in future datasets.

The poor predictive performance of the MIC classifier reinforces the finding that broad taxonomic groupings do not adequately capture MIC-level variation among the tested organisms. Variation in outer membrane permeability and efflux pump expression among Gram-negative species can produce substantially different MIC values even within the same broad taxonomic class (Nikaido, 1998). Species-specific biological characteristics are therefore likely to influence MIC outcomes more strongly than broad taxonomic classification alone. In the present study, this finding indicates that coarse categorical predictors were insufficient for the available MIC outcomes.

The use of Random Forest for antimicrobial discovery is consistent with previous applications of the algorithm to natural-product and antimicrobial datasets. Random Forest models have been used to predict active antimicrobial compounds from chemical descriptor data (Jana et al., 2024) and to identify potential natural antibiotic plants from traditional herbal formulations (Nasution et al., 2022). The present study extends this computational approach to mushroom-derived extracts, but the findings should remain within the scope of the current dataset and should not be interpreted as evidence of broad cross-system generalizability.

From a methodological perspective, LOOCV was selected as the primary evaluation strategy because it maximized the use of the limited observations. However, LOOCV alone does not fully address the clustering introduced by repeated batch-level predictors. The supplementary leave-one-batch-out analysis was therefore conducted specifically to assess whether this clustering inflated the reported performance and did not indicate materially lower performance. Nevertheless, with only three cultivation batches available, neither cross-validation scheme can substitute for validation on independent, larger datasets, and the reported performance metrics should be interpreted cautiously.

Overall, these findings support the feasibility of integrating machine learning into antimicrobial screening workflows for mushroom-derived natural products, consistent with the broader application of ML to natural-product bioactivity prediction (Yuan et al., 2023). Rather than replacing laboratory experiments, predictive models may serve as decision-support tools for prioritizing extracts or experimental conditions for subsequent validation. Broader applicability will require substantially larger datasets incorporating multiple substrates, mushroom species, and replicate-resolved biochemical measurements before robust, generalizable predictive models can be developed.

## 5. Conclusion

This proof-of-concept study demonstrated the feasibility of applying machine learning to experimentally generated antimicrobial assay data from *Pleurotus ostreatus* cultivated on cassava peel. The Random Forest regression model showed moderate internal predictive performance for zone of inhibition, indicating that computational modelling can identify preliminary predictor–response associations within a small experimental dataset. Extraction solvent was the most influential predictor, while the batch-level mycochemical variables contributed minimally to model importance. These findings are consistent with substrate-driven bioactivity patterns reported in the parent study (Adetuwo et al., 2026).

In contrast, the MIC classification model showed limited predictive capability, indicating that the broad pathogen taxonomic groupings and available predictors were insufficient to reliably discriminate the two observed MIC groups. Future work should incorporate larger, multi-substrate and multi-species datasets with replicate-resolved biochemical profiling to determine whether the relationships identified here persist across broader experimental conditions and to support development of more generalizable predictive models.

## Software and Data Availability

All machine learning analyses were implemented in Python 3 using the scikit-learn library (Pedregosa et al., 2011) for Random Forest regression, Random Forest classification, and leave-one-out cross-validation. Data visualization was performed using Matplotlib. The raw antimicrobial assay dataset (zone of inhibition and minimum inhibitory concentration values, with associated batch-level mycochemical measurements) and the analysis scripts used to generate the reported models and figures have been deposited in Zenodo and are publicly available at https://doi.org/10.5281/zenodo.21848489 (Adetuwo, 2026). The repository is organized into raw data (one file per assay type, matching the original laboratory notebook records), a processed/merged analysis-ready dataset, numbered scripts covering the full pipeline from raw data to figures, and the resulting model outputs and figures; a data dictionary and README accompanying the deposit describe every column and file in full.

